# Probing Neural Networks for Dynamic Switches of Communication Pathways

**DOI:** 10.1101/523522

**Authors:** Holger Finger, Richard Gast, Christian Gerloff, Andreas K. Engel, Peter König

## Abstract

Dynamic communication and routing play important roles in the human brain in order to facilitate flexibility in task solving and thought processes. Here, we present a network perturbation methodology that allows investigating dynamic switching between different network pathways based on phase offsets between two external oscillatory drivers. We apply this method in a computational model of the human connectome with delay-coupled neural masses. To analyze dynamic switching of pathways, we define four new metrics that measure dynamic network response properties for pairs of stimulated nodes. Evaluating these metrics for all network pathways, we found a broad spectrum of pathways with distinct dynamic properties and switching behaviors. We show that network pathways can have characteristic timescales and thus specific preferences for the phase lag between the regions they connect. Specifically, we identified pairs of network nodes whose connecting paths can either be (1) insensitive to the phase relationship between the node pair, (2) turned on and off via changes in the phase relationship between the node pair, or (3) switched between via changes in the phase relationship between the node pair. Regarding the latter, we found that 33% of node pairs can switch their communication from one pathway to another depending on their phase offsets. This reveals a potential mechanistic role that phase offsets and coupling delays might play for the dynamic information routing via communication pathways in the brain.

**Author summary:** A big challenge in elucidating information processing in the brain is to understand the neural mechanisms that dynamically organize the communication between different brain regions in a flexible and task-dependent manner. In this theoretical study, we present an approach to investigate the routing and gating of information flow along different pathways from one region to another. We show that stimulation of the brain at two sites with different frequencies and oscillatory phases can reveal the underlying effective connectivity. This yields new insights into the underlying processes that govern dynamic switches in the communication pathways between remote sites of the brain.

## Introduction

Over the past decades it has been shown that the brain, facing a specific task or not, exhibits well-structured functional connectivity [1–5]. This has been specifically investigated for resting-state networks [6–9], but also for other networks when the brain is performing different tasks [8, 10]. These findings lead to the idea that networks of dynamically coupled neural populations reflect an inherent functional organization of the brain which is optimized to perform a wide range of tasks it encounters frequently [11, 12]. If faced with a task that requires synchronization between brain areas not typically coupled at rest, this organization has to be altered temporarily in order to perform that task efficiently [13–15].

One mechanism that has been proposed for such dynamic re-organization is top-down suppression of distractor areas (or task-irrelevant areas) by a slow, oscillatory rhythm compared to the neural rhythm at which the signal is being processed [16–18]. While this suppression mechanism acts on individual nodes in the network, dynamic re-organization by changes in the phase relationships could act directly on the edges of the network by modulating the effective connectivity between nodes [19–21]. Here, we focus on the latter and consider the dynamic re-organization as a self-organizing property of the network that flexibly adapts functional coupling to changes in extrinsic stimulation. According to the communication-through-coherence theory [19, 20], the phase relationship between two neural oscillators affects their success in communication. A neural population which tries to establish communication with other target populations at a certain frequency is more successful if the signals arrive at the excitable phase of the target populations. In other words, successful communication between two neural oscillators is reflected by their amount of entrainment. The functional coupling pattern that arises from this process could be temporarily altered by advancing or slowing down the phase of each of the neural oscillators. This way, changes in phase relationships between brain areas could serve as a mechanism for the re-organization of effective connectivity.

Within a complex network like the human brain, multiple structural pathways exist between most pairs of nodes given a sufficiently high spatial resolution. Since functional coupling patterns are established via synchronization along such pathways, we set out to identify general principles of how these pathways interact with each other during synchronization processes. Several studies have placed emphasis on the importance of information transmission delay for synchronization processes as well as its role in the formation of functional clusters in the brain [5, 20, 22–32]. In their fundamental work on the dynamic implications of discrete delays in networks of rate-based neurons, Roxin and colleagues were able to demonstrate that the incorporation of delays enriches the dynamic repertoire of the system, allowing for various periodic and aperiodic behaviors [33, 34]. Using connectome models composed of delay-coupled Kuramoto oscillators, Petkoski and colleagues showed that delay distributions in heterogeneously coupled networks can influence whether nodes tend to synchronize in phase or anti-phase [32]. Furthermore, Gosh and colleagues showed in a connectome model of coupled Fitz-Hugh Nagumo systems that the incorporation of realistic network delays allowed the noise-driven model to show a dynamic repertoire similar to experimentally reported dynamics of resting-state functional connectivities [35]. Here, we investigate the interplay between coupling delays and synchronization phase offsets in a model of the human connectome. In the case of two delay-coupled neural oscillators, a tight relationship between their optimal phase relationship and coupling delay can be expected [20]. Given a complex network with multiple communication pathways between node pairs, this raises the question of whether the set of active communication pathways depends on the phase offset between the nodes. Finding a difference in this set of utilized communication pathways over varying phase offset strongly implies that communication pathways express preferences for the phase relationship between the nodes they connect. These preferences will be influenced by the time delays of the communication pathways, which are determined by axonal signal transmission delays as well as rise and decay properties of the post-synaptic response. Here, we focus on the former and assume the latter to be relatively homogeneous across the cortico-cortical connections considered in our model, since they all resemble glutamatergic synaptic connections. In particular, we predict that two brain regions trying to communicate at a certain frequency with a given phase offset will use only a fraction of their available communication paths. Furthermore, we predict the selection of communication paths to be influenced by their interaction time delays and the phase offset at which the communication is initially attempted.

To test these hypotheses, we investigate cortico-cortical communication processes via a computational whole-brain network model of delay-coupled, oscillating nodes. We introduce a set-up with two oscillatory drivers to probe the network for changes in interactions between the stimulated pair of nodes. Importantly, our stimulation approach relies on the entrainment of the given pair of nodes to oscillate at the same frequency, but with a certain phase lag relative to each other. While Fig 1A illustrates the overall extrinsic stimulation setup, Fig 1B visualizes why it is necessary to stimulate at different phase lags between the stimulation signals. In particular, Fig 1B shows that even in the absence of any interaction through the network, there is the externally induced trivial coherence between the two stimulated nodes. Thus, the coherence is measured for many different stimulation phase offsets and the measurement with the lowest coherence is chosen as the baseline. Any deviation in the coherence from this baseline can be attributed to induced changes in the coupling between the two stimulated nodes through the network. As indicated in Fig 1C and 1D, this may happen due to a switching in the communication pathways. Comparing the coherence along different pathways over different stimulation phase offsets then reveals the phase preferences for different routes. We speculate that these differences in phase preferences at different pathways could act as a switching and gating mechanism used by the brain to establish communication between remote brain areas when needed. Our method allows the investigation of these mechanisms by probing the network for these dynamic switches in communication pathways.

**Fig 1.**
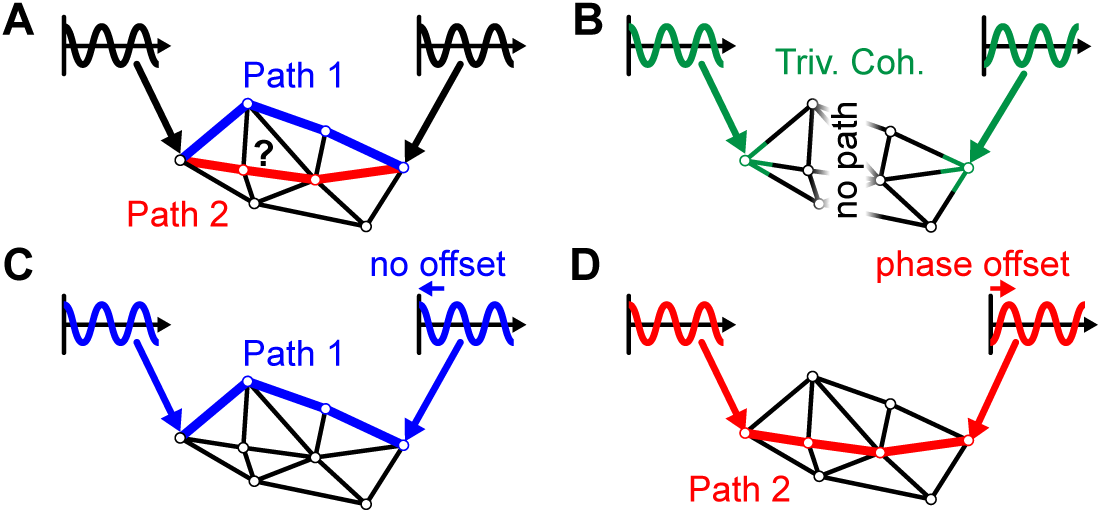
Illustration of the stimulation methodology and different possible outcomes. (A) The stimulation at the two nodes could potentially activate different paths (blue or red) in the network. (B) The simultaneous stimulation at the two nodes could induce a trivial coherence (green) between the two nodes even in the absence of any interaction between the two nodes through the network. (C) In this example a stimulation with no phase offset induces a network interaction between the two stimulated nodes through path 1 (blue). (D) In contrast, when stimulating with a phase offset of *π* another path is activated (red).

## Computational Model

Our computational model (Fig 2) is based on the widely used Jansen-Rit neural mass model [36] which employs a mean-field approach to model the interaction between a pyramidal cell population (green), an excitatory interneuron population (blue), and an inhibitory interneuron population (red), as illustrated with the relevant equations in the Figure. The function *σ*(*V*) transforms the average membrane potentials to average firing rates via a parameterized sigmoid (depicted in cyan). The standard parametrization originally proposed by Jansen and Rit reflects cortical oscillatory activity in the alpha frequency band. These parameters were chosen based on experimental findings in the neuroscience literature and are reported in the Materials and Methods section. Since the aim of this article is the investigation of the effect of pathway time delays on neural synchronization processes and not the effect of information processing delays at the node level (arising from time constants of cell membranes, synapses etc.), we decided to use this standard parametrization for each node in our network [36]. For the purpose of investigating networks of extrinsically perturbed Jansen-Rit models, the following two extensions were added: First, we coupled multiple Jansen-Rit nodes via delayed, weighted connections between their pyramidal cell populations (depicted in yellow). In our model this pyramidal cell population is interpreted as a lumped representation of all projection cells within a modeled brain area, whereas the excitatory and inhibitory interneuron populations describe local feedback loops of that pyramidal cell population. We avoid the original layer-specific (infragranular, granular, and supragranular) interpretation offered by Jansen and Rit [36] because inter-areal connectivities would be subject to layer-specific profiles [37] which would be hard to estimate reliably for a whole-brain network model. Note that this approach is in accordance with previous simulation studies of networks of multiple coupled Jansen-Rit models that employed pyramidal to pyramidal cell coupling as well [38–42]. Secondly, weak external drivers were applied at two stimulation sites influencing the average membrane potential of the pyramidal cells with phase offset Δ*φ* between the two drivers (depicted in purple). By applying the stimulation to the pyramidal cells, it enters at the same population which also receives the network input. It can, thus, either be conceived as input of a neural origin not explicitly incorporated in our model or as an extrinsic input that mainly affects the excitability of the pyramidal cells.

**Fig 2.**
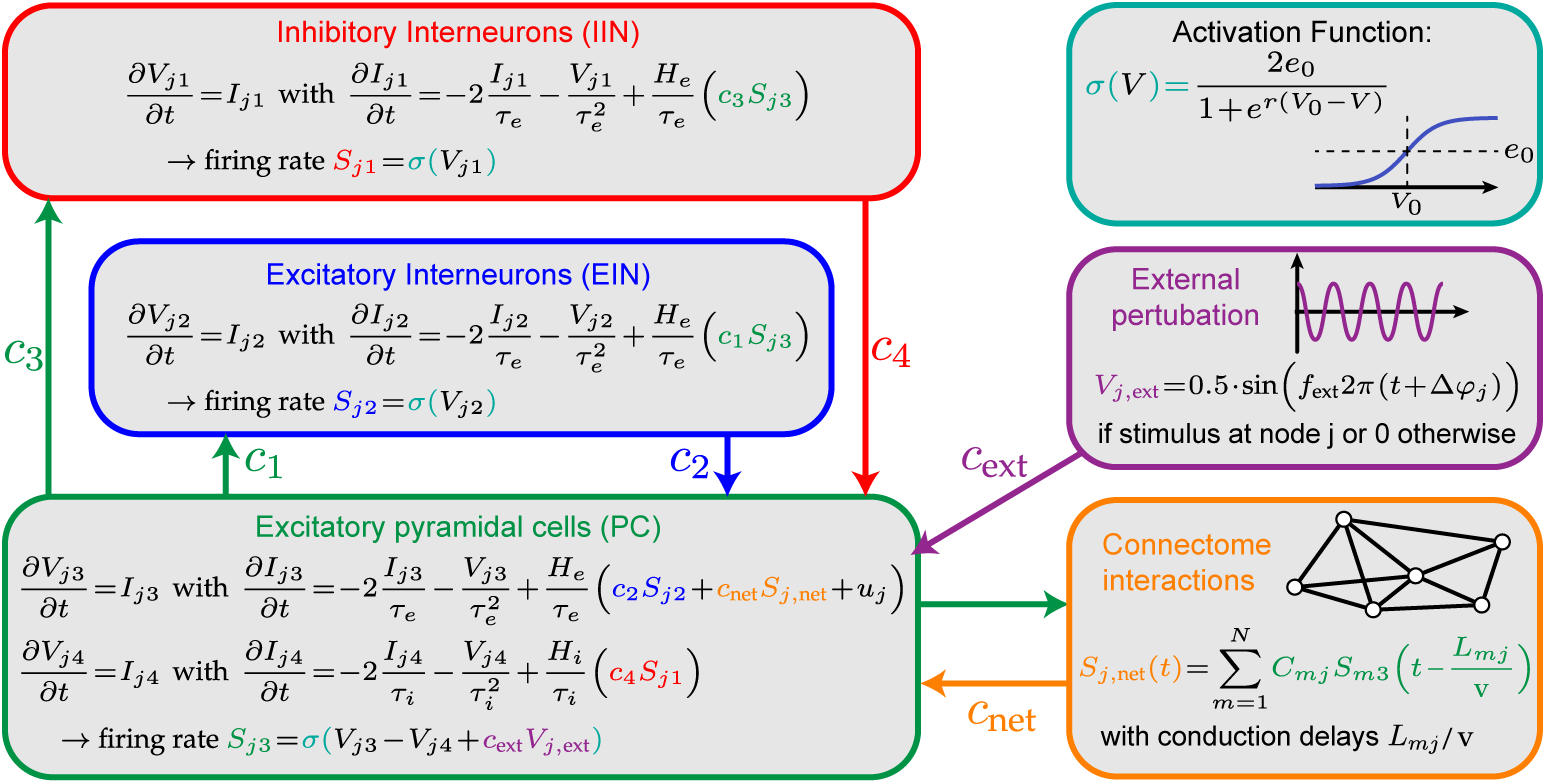
Neural mass model with external stimulation. A schematic of the neural mass model showing the interactions between the three neuronal populations (pyramidal cells, excitatory interneurons, inhibitory interneurons). Each post-synaptic potential is modeled using two differential equations (average membrane potentials *V* and average synaptic currents *I*. Several of these neural mass models are interacting through a connectivity matrix (yellow). The external perturbation (purple) modulates the average membrane potential of pyramidal cells at 2 nodes in the network.

## Results

We will first present our results of the simulations in simple networks of only 2 or 3 nodes. Specifically, we show how the coherence between a pair of stimulated nodes depends on the transmission delay of their connections and on the relative phase offsets between the two stimulation signals. Subsequently, we move on to a model of the cortico-cortical synchronization processes within a single hemisphere of the human brain. Cortico-cortical coupling strengths and delays were informed by an approximation of the human connectome obtained from diffusion tensor imaging (DTI) data. Additionally, we fit the model to match functional connectivities calculated from electroencephalography (EEG) recordings. In this model, we show how the coherence between stimulated nodes changes over phase lags and how this effect relates to the connectedness and distance of the nodes. In a final step, we identify the pathways responsible for the interaction between the stimulated nodes, analyze their phase lag preferences and identify cases of phase-related switching between pathways. For all simulations, we use the computational model defined in the previous section.

### Simple Models With 2 or 3 Nodes

The idea behind the extrinsic stimulation approach can be well explained using a simple toy-model of 2 directly coupled Jansen-Rit nodes, where each node is stimulated with a *f*_ext_ = 11 Hz sinusoidal signal with strength *c*_ext_ = 0.1 mV. Fig 3 shows the coherence between the driven nodes for systematic changes in the phase offset between the stimuli and the distance between the coupled nodes. While uni-directionally coupled nodes can have preferences for any stimulation phase offset, as shown in Fig 3A, bi-directionally coupled nodes are more susceptible for stimulation at in- or anti-phase (see Fig 3B). The latter is in line with other studies that showed neural synchronization of remote, delay-connected neurons or neural populations to typically appear at in- or anti-phase [31, 32, 43]. This shows that the communication between coupled pairs of nodes can be modulated by stimulation and that communication pathways can have characteristic stimulation phase offset preferences, depending on their length [44].

**Fig 3.**
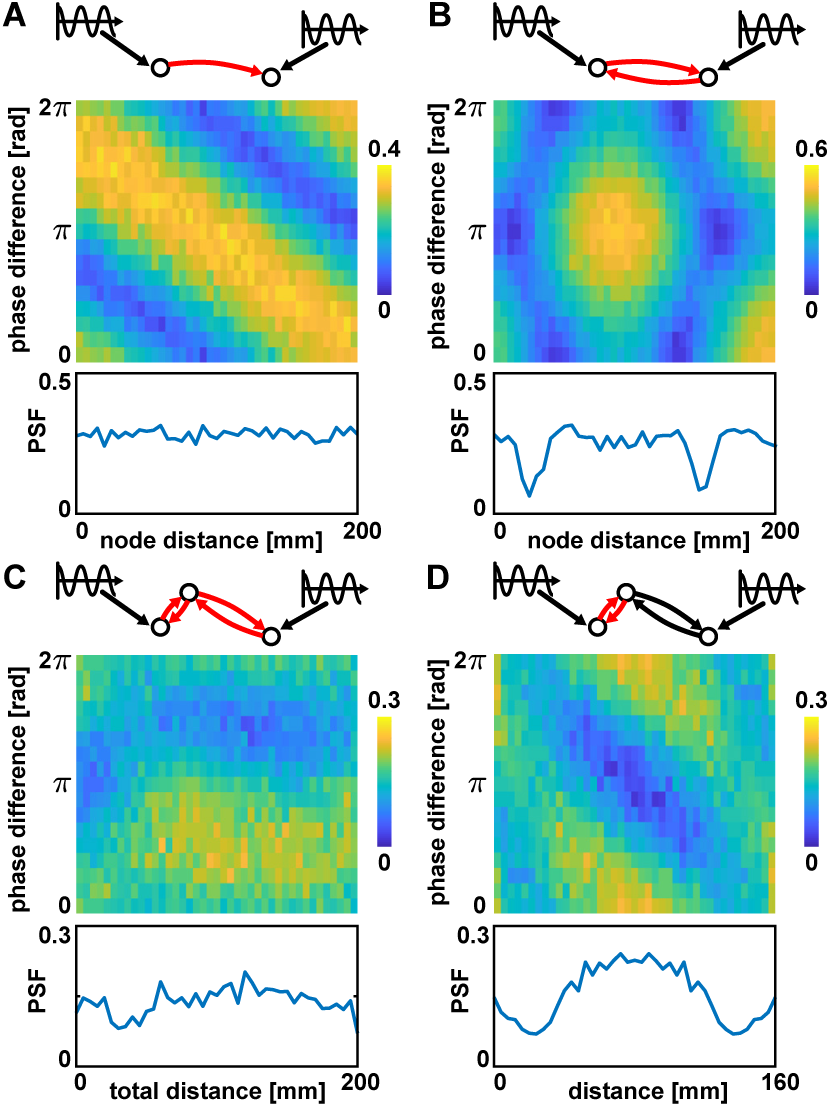
Simulations with 2 or 3 nodes. Red edges indicate the connections for which the distance was varied. Color indicates the coherence between the two driven nodes. The pathway synchronization facilitation (PSF) values are shown at the bottom of each panel. (A) Nodes with direct uni-directional coupling. (B) Nodes with direct bi-directional coupling. (C) Nodes with indirect (via a third node) bi-directional couplings. The intermediate node was placed at 25 % of the total connection distance, while the overall distance between the outer nodes was varied. (D) Nodes with indirect bi-directional couplings where the overall distance was kept at 160 mm, while the intermediate node was positioned at varying positions along the connection. Parameters used in all panels: *v* = 2.6 m/s, *c*_net_ = 14, and *C*_*mj*_ = 0.1 if there is a connection from node m to node j or 0 otherwise.

To quantify the modulation of communication, we define the pathway-synchronization-facilitation (PSF), measuring for a given pair of weakly stimulated network nodes *k*_*i*_ and *k*_*j*_ how their interaction is dependent on specific phase offsets:

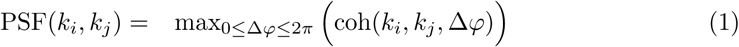

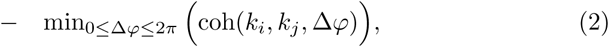

where coh(*k*_*i*_, *k*_*j*_, Δ*φ*) is the coherence between network nodes *k*_*i*_ and *k*_*j*_ for stimulation phase offset Δ*φ*. The PSF is high for node pairs if their coherence is high for one stimulation phase offset and low for another, i.e., the relative phase of the stimulation at the two sites matters strongly. The PSF curves in Fig 3A and 3B show that in both cases there is a PSF effect (PSF > 0) and in the case of bi-directionally coupled nodes the strength of this effect depends on the distance between the nodes.

To extend this idea to communication via indirect pathways, we investigated synchronization between 2 nodes connected only indirectly via a third intermediate node. We used bi-directional couplings for both connections and both end nodes were stimulated as described previously. As can be seen in Fig 3C and 3D, the interaction between the two weakly stimulated nodes not only depended on the length of the connection, but mostly on the relative position of the third node on the indirect path. Thus, phase lag preferences of pathways connecting two target nodes arise primarily from the distance between intermittent nodes along the pathway. In a heterogeneous network like the connectome, we hence expect to find node pairs connected by pathways with distinct phase lag preferences. Depending on the stimulation phase offset at the target nodes, we expect to be able to induce a switch in the pathway the nodes employ to entrain each other.

### Connectome Model Without Stimulation

As shown in the previous section, the coherence in a network of only three nodes can already exhibit very complex dependencies on the stimulation phase offset. To extend our analysis to network communication patterns in the case of a complex network with multiple competing pathways, we used a model of 33 delay-connected nodes, representing one hemisphere of the human connectome [30]. The structural connectivity matrix was obtained from DTI-based tractography data as described in the Materials and Methods section. Fig 4A shows the sparse connectivity matrix *C*_*mj*_ used to connect the 33 regions and Fig 4B the corresponding average fiber lengths *L*_*mj*_ between region pairs. These two matrices are used for connecting the different populations of excitatory pyramidal cells (green in Fig 2) in a delay-coupled network of a single hemisphere.

**Fig 4.**
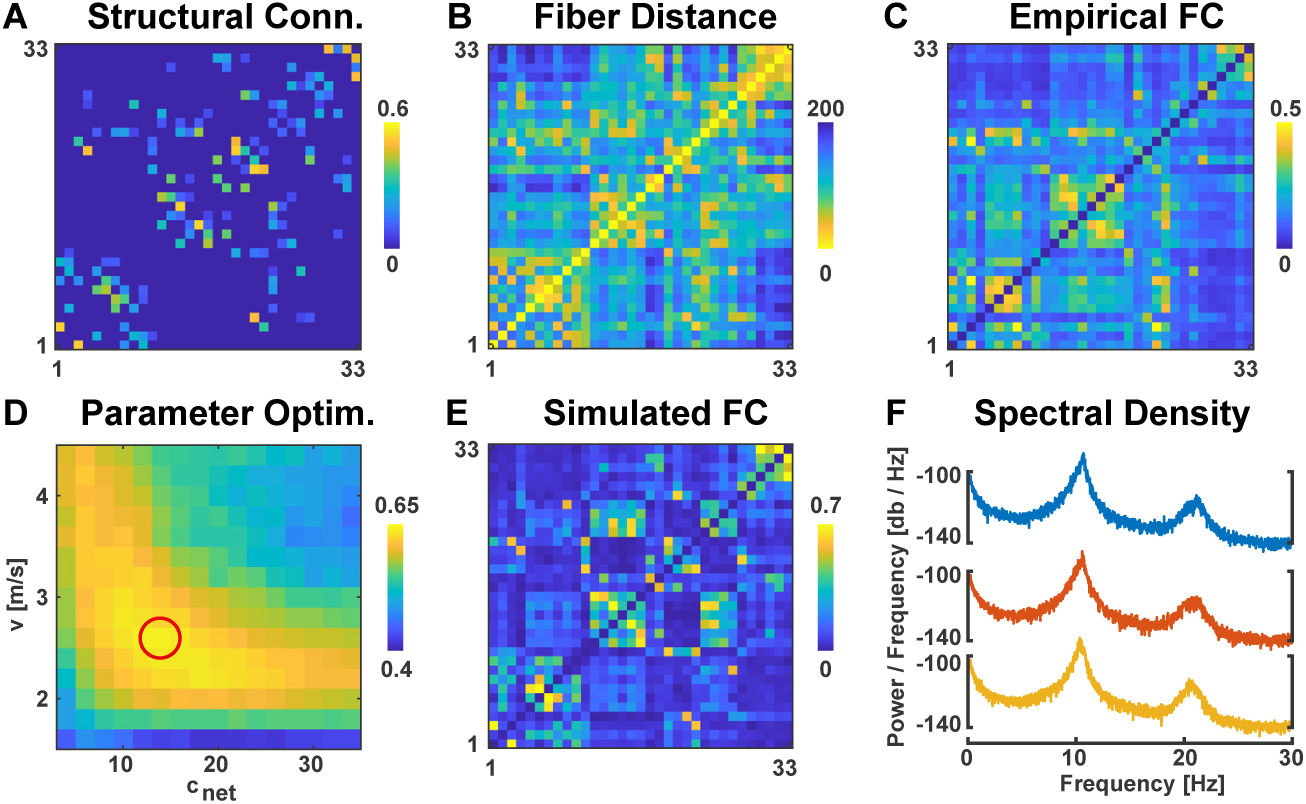
Pairwise measures of connectivity and distance between all 33×33 region pairs. (A) Structural connectivity matrix *C*_*mj*_ with all connections smaller than 0.1 set to 0. (B) Inter-regional average fiber distances *L*_*mj*_ in mm. (C) Functional connectivity matrix derived from coherence of EEG data bandpass-filtered around 10 Hz. (D) Correlation between the empirical and the simulated functional connectivity matrices for different global velocities *v* and global connection strengths *c*_*net*_. Encircled in red is the parameter set chosen for all subsequent connectome model simulations. (E) Functional connectivity derived from coherence of neural mass model simulations bandpass-filtered around 10 Hz. (F) Power-spectral densities of 3 representative network nodes.

For the same 33 regions, EEG resting-state recordings from the same subjects were used to calculate pairwise coherences in the 10 Hz range as shown in Fig 4C. To resemble this empirical resting-state functional connectivity, we used a grid-search over the two remaining free parameters that were not defined in [36]: the global scaling of the connection weights *c*_*net*_ and the global velocity *v* that is used to scale the distance data resulting in pair-wise signal transmission delays. We simulated the 33 connected neural-mass-models and processed the time-series of the pyramidal cells in the same way as the EEG data, i.e. we band-pass filtered the time series at 10 Hz, applied the Hilbert transform, and calculated the pair-wise coherence. This yielded a 33 x 33 functional connectivity matrix which we compared to the empirical functional connectivity by calculating the Pearson correlation coefficient.

The selection of parameters was based on the rationale to match the functional connectivity observed in the network model as well as possible to empirical EEG-based functional connectivity from human subjects. By fitting the velocity, we ensured that our pathway delays reflect realistic, empirically observed timescales of cortico-cortical interactions (Fig 4D). We selected the parameters *c*_*net*_ = 14 and *v* = 2.6 m/s for subsequent analyses which show a high correlation to the empirical functional connectivity (r = 0.64, p < .0001). Notice that this correlation is comparable with values of other bottom-up models reported in the literature [30, 45], which is remarkable considering that we set a substantial amount of structural connections to 0. The simulated functional connectivity using these selected parameters is shown in Fig 4E. A comparison of these different connectivity matrices (Fig 4A,B,C,E) revealed that the empirical functional connectivity correlated more strongly with the inter-regional fiber distances (*r*_*CB*_ = −0.70) than with the structural connectivity (*r*_*CA*_ = 0.55), while the simulated functional connectivity correlated more strongly with structural connectivity (*r*_*EA*_ = 0.78) than with the inter-regional fiber distances (*r*_*EB*_ = −0.71).

As a last analysis of the unperturbed network, we evaluated the power spectral density of each network node (Fig 4F). All 33 nodes showed a frequency peak around 11 Hz.

### Connectome Model With Stimulation

Based on this model of cortical activity, we used the stimulation approach to investigate how pathways facilitate synchronization between network nodes at certain phase lags between the nodes. Specifically, we weakly stimulated different pairs of cortical regions with varying phase offsets between the two stimulation signals while measuring the coherence between the stimulated nodes at each phase offset. As argued above, finding differences in the coherence over stimulation phase offsets would indicate phase-specific communication modulation between the stimulated nodes. Before analyzing PSF effects in the connectome model, it was necessary to determine the optimal stimulation frequency and strength for this model, because complex synchronization patterns can arise in networks of delay-coupled non-linear elements [33], especially when perturbed extrinsically [41]. This was performed in two steps. First, we determined the optimal frequency by extrinsic stimulation of a single network node. Second, we determined the optimal stimulation strength by stimulation of two network nodes at that frequency.

Since our main analysis will focus on coupling effects through different network paths between two stimulated nodes, we aimed to find the stimulation frequency at which the extrinsic stimulation has a high penetration depth into the network (as measured by the mean coherence to all network nodes). Notice that the stimulation frequency with the optimal network penetration depth may not necessarily coincide with the optimal stimulation frequency to entrain the stimulated network node. To determine the optimal stimulation frequency, we stimulated a single region in our network with a stimulus of varying frequency (4-30 Hz) and strength (0.03-4 mV) while evaluating the mean coherence between the stimulus and all nodes in the network. As Fig 5A shows, this average coherence to the full network shows a strong peak at 11 Hz, which is the intrinsic frequency of the unperturbed network (see Fig 4F). The mean coherence of the stimulation signal to the stimulated node and its direct neighbours can be observed in Fig 5B, while the coherence to only the stimulated node can be observed in Fig 5C (always averaged over simulations in which the extrinsic stimulation was applied to each of the 33 regions). The latter reflects the well-studied relationship between a driver and an oscillator described by the so-called Arnold tongue [46], except that this coherence to the stimulated node was higher at driving frequencies that are different to the intrinsic network frequency of 11 Hz (see Fig 5C). This reflects the competition between entrainment via the extrinsic stimulation and via the network. At the intrinsic frequency of the network (11 Hz) and its harmonics, the driver has to compete with the network in entraining the stimulated node at a certain phase relationship. This competition decays with the deviation of the driving frequency from the networks’ intrinsic frequency and its harmonics. Thus, the Arnold tongue of a single driven node embedded in the connectome model shows coherence peaks shifted away from its intrinsic frequency. However, as can be seen in Fig 5B, entrainment of the direct neighbours of the stimulated node is still strongest at the networks’ intrinsic frequency, which explains the mismatch between the optimal stimulation frequency for penetrating the network (Fig 5A) vs. entraining a single node (Fig 5C). To ensure that the network is not pushed into a fully synchronized state by the extrinsic perturbation, we also evaluated the Kuramoto order parameter of the network (see Fig 5D). This metric reflects the level of synchronization in the network, with a value of 1 implying full synchronization and a value of 0 implying a fully asynchronous state. As can be seen in Fig 5D, the Kuramoto order parameter varies between 0.1 and 0.25 across extrinsic stimulation parametrizations and expresses a relatively low value in this range for an 11 Hz driver. Based on these results, we set the frequency of our stimulus to 11 Hz, at which the network (and not only the directly stimulated node) was most susceptible for entrainment by an external stimulation, but was not driven into a fully synchronized state.

**Fig 5.**
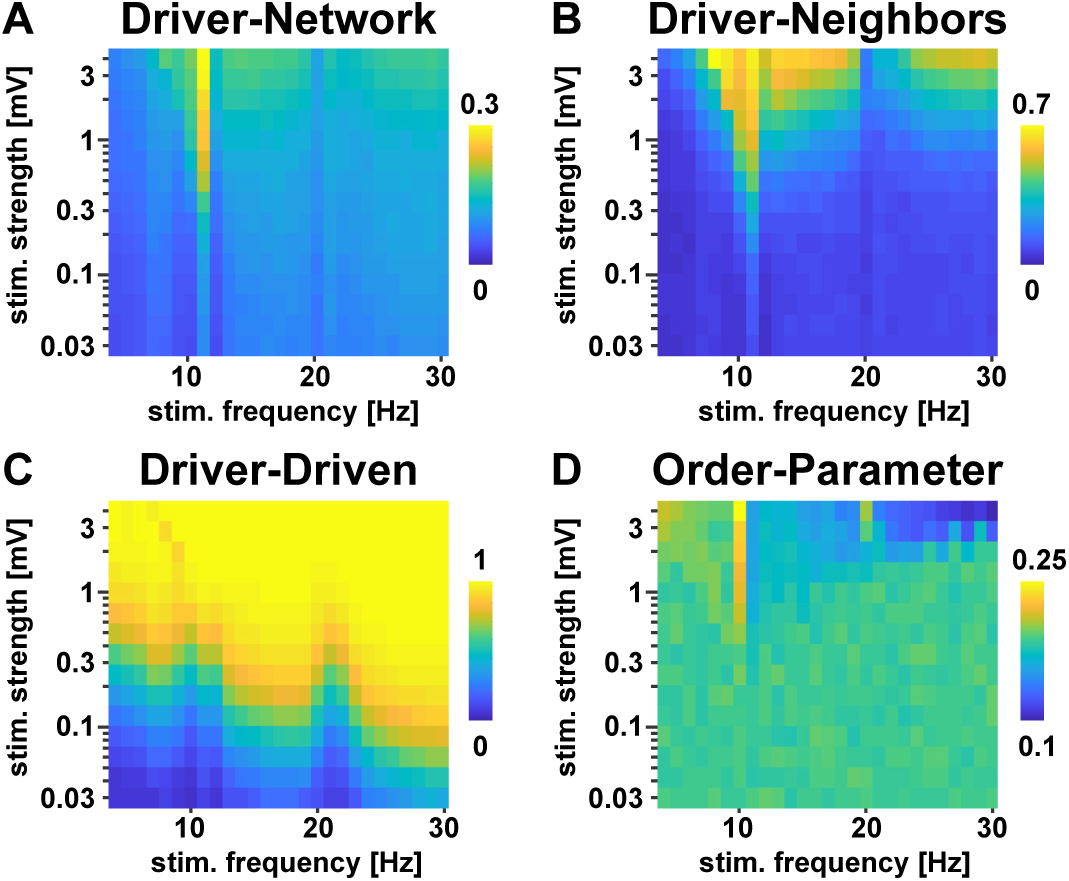
Stimulation parameter evaluation in the connectome with a single driver. All panels show averaged results that were obtained by first placing the stimulus at each of the 33 regions individually and finally averaging over these different stimulus locations. (A) Average coherence between the stimulus and the full network for varying stimulus strength and frequency. Each region was stimulated and the average coherence between the stimulation signal and each network node was calculated. (B) Average coherence between the stimulus and all direct neighbors of the stimulated region. (C) Coherence between the stimulus and the stimulated region. (D) Kuramoto order parameter of the network for each stimulation strength and frequency.

In a second step, we stimulated pairs of nodes with 11 Hz stimuli. We varied the stimulus strength (0.25 - 1 mV) and the relative phase offset between the stimuli (0 − 2*π*) while evaluating the coherence between the stimulated nodes. All other parameters were chosen to be the same as for the previous simulation. The variability in the coherence between stimulated region pairs that we observed over stimulation phase offsets (as depicted for 2 example region pairs in Fig 6A and 6B) shows that the stimulated region pairs interacted with each other and that the interactions expressed a characteristic profile of phase offset preferences. As can be seen in Fig 6A and 6B, the variance of the coherence over phase offsets depended on the stimulation strength. Based on visual inspection of the coherence patterns of 20 different region pairs, we chose our stimulus scaling to be *c*_ext_ = 0.5 mV, leaving the variance of the post-synaptic potential of the neural masses in a biologically plausible range and such that the external driver is relatively weak in comparison to the internal network dynamics (the membrane potential fluctuations of a single Jansen-Rit node are in the order of 1-10 mV). This gave us the final set of global model parameters which were used throughout all subsequent simulations and are reported in Table 1.

**Table 1.** Optimized Model Parameters.

| Param. | Value | Interpretation |
| --- | --- | --- |
| $c_{\text{net}}$ | 14 | global connection strength scaling |
| $v$ | 2.6 m/s | global velocity scaling |
| $f_{\text{ext}}$ | 11 Hz | extrinsic stimulation frequency |
| $c_{\text{ext}}$ | 500 $\mu\text{V}$ | amplitude of extrinsic stimuli |

**Fig 6.**
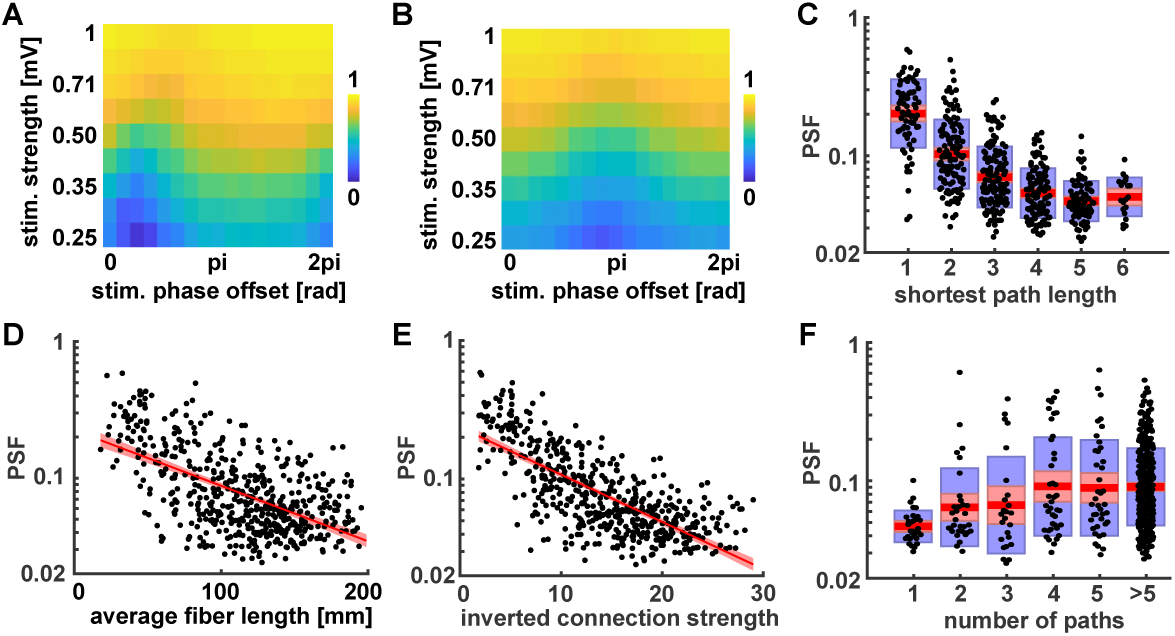
Pairwise stimulation with different phase offsets in the connectome. (A & B) Coherence between two stimulated nodes for varying stimulation strength and phase offset. A and B correspond to two different example node pairs that were stimulated to demonstrate the variability in phase lag preferences expressed by different node pairs. Panels (C-F) show the Pathway Synchronization Facilitation (PSF) on a logarithmic vertical axis with red areas indicating 95 % confidence intervals and blue boxes indicating 1 standard deviation. (C) Dependence of PSF effect on shortest path length between the node pairs. (D) Correlation between PSF effect and summed connection lengths along the shortest pathway between the stimulated nodes (calculated from matrix *L*_*mj*_). (E) Correlation between PSF effect and the inverted connection strength between the stimulated nodes, which is measured as the Dijkstra distance based on the inverted connection strengths 1*/C*_*mj*_. (F) Dependence of the PSF effect on the number of paths connecting the stimulated nodes.

To statistically confirm the variance in the coherence between stimulated region pairs over phase offsets, we ran simulations with subsequent stimulations of each possible node pair in the network. Again, we varied the phase offset between the two stimuli (16 equally spaced phase offsets between 0 and 2*π*) and evaluated the coherence between the stimulated nodes for each phase offset. Subsequently, those coherences were used to calculate the PSF for each region pair as defined in Eq (1). Using a one-sample t-test, we found the PSF effect to be significantly larger than zero (mean = 0.10021, CI=[0.092875, 0.10755], t = 26.8268, p < .0001). Hence, we were able to show with our extrinsic stimulation approach that pathways facilitated synchronization between cortical nodes and that the facilitatory strength depended on the phase lag between the region’s average PSPs.

With the PSF effect established, we continued by investigating its dependence on certain features of the underlying structural connectivity graph. For this purpose, we searched for all possible pathways between each pair of stimulated nodes based on the structural connectivity matrix reported in Fig 4A. Since every stimulated pair of nodes was connected by at least one path via at most 6 edges, we restricted the search to pathways including 6 edges at maximum. With these pathways at hand, we started out by evaluating how the PSF effect changed with increasing network distance. An analysis of variance showed that the effect of shortest path length (minimum number of edges seperating a pair) on log(PSF) was significant, F(5,521) = 75.22, p < .0001. As can be seen in Fig 6C, we observed the trend that the PSF effect decreases with the number of nodes separating the stimulated nodes. Furthermore, as depicted in Fig 6D, this trend was supported by a significant correlation between the PSF effect and the length of the shortest pathway between the stimulated nodes (r = -0.57, p < .0001), a measurement that is strongly related to both interregional distance and minimal number of separating edges. Thus, we conclude that there is a tendency for a decrease in the interaction of stimulated node pairs with increasing network distance, which can be measured either as the number of interposed edges or as the summed up length of the edges of the shortest pathway connecting the nodes.

Next, we investigated the dependence of the PSF effect on the connectedness between the stimulated nodes. In this regard, we characterize the connectedness either by the connection strength along the shortest path (evaluated in Fig 6E) or by the number of different pathways connecting the two nodes (evaluated in Fig 6F). In the first case, we found a high correlation between the Dijkstra distance, calculated from the inverted connection strengths 1*/C*_*mj*_, and the PSF (r = -0.73, p < .0001) which is depicted in Fig 6E. This confirms previous findings of the dependence of the (directed) interactions between delay-coupled oscillators on their connection strength [29, 32].

In the second case (Fig 6F), we measure the connectedness by the number of different pathways connecting the two nodes. It was necessary to limit this analysis to include only pathways with up to 6 edges, because of the combinatorial growth of the number of possible pathways with more edges. In other words, considering pathways with more edges could lead to scenarios in which node pairs show a large number of connected paths, even though those pathways all have to pass through many intermediate nodes and might not contribute to the interaction of these nodes through the network at all. Thus, to render this measure of connectedness less noisy, we decided to exclude paths with more than 6 edges. An analysis of variance showed that the effect of the number of connecting paths (only counting paths with 5 edges or less, and all nodes with more than 5 connecting paths were pooled into one level) on log(PSF) was significant, F(5,501) = 7.42, p < .0001. Together, these results (Fig 6 and Fig 3) reveal how the variance of the interaction of network nodes over different phase lags (as measured by the PFS) depends both on the strength and delay of their connecting pathways.

### Evaluation of Pathway Activation

To identify, how different pathways between stimulated node pairs contribute to the PSF effect, we next aimed to quantify how much a particular pathway was involved in the node interaction at a given phase offset. For this analysis, we define the pathway activation (PA) for a pathway through n nodes *k*_*i*_ with *i* = 1..*n* at a phase offset Δ*φ* as the product of the pairwise coherences between neighboring pathway nodes:

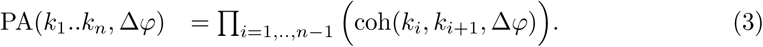

In other words, if communication fails at any point along a pathway, leading to a reduced coherence between the involved nodes, this is considered to be a bottleneck for the information flowing through that pathway. This is similar to the definition of efficiency of neural communication via communication pathways provided in [21]. Thus, for the remainder of this article, we will use changes in information flow and changes in pathway activation interchangeably. Furthermore, this metric ensures that short pathways are preferred over longer pathways, since the coherence is bound to the interval between 0 and 1.

We evaluated the pathway activation (PA) for all pathways of up to *n* = 5 nodes connecting a given pair of stimulated nodes for all stimulation phase offsets. Doing this for each stimulated node pair, we found different classes of pathway interactions: Some pairs show only a very small selectivity for the stimulation phase offset (Fig 7A), while other node pairs were connected by paths with PA values with a strong dependence on the phase offset (Fig 7B, 7C). Moreover, some of these node pairs switched their interaction between different pathways depending on the stimulation phase offset. This is shown in Fig 7C, while the pathways between which the switching occurs are visualized in Fig 7D and 7E.

**Fig 7.**
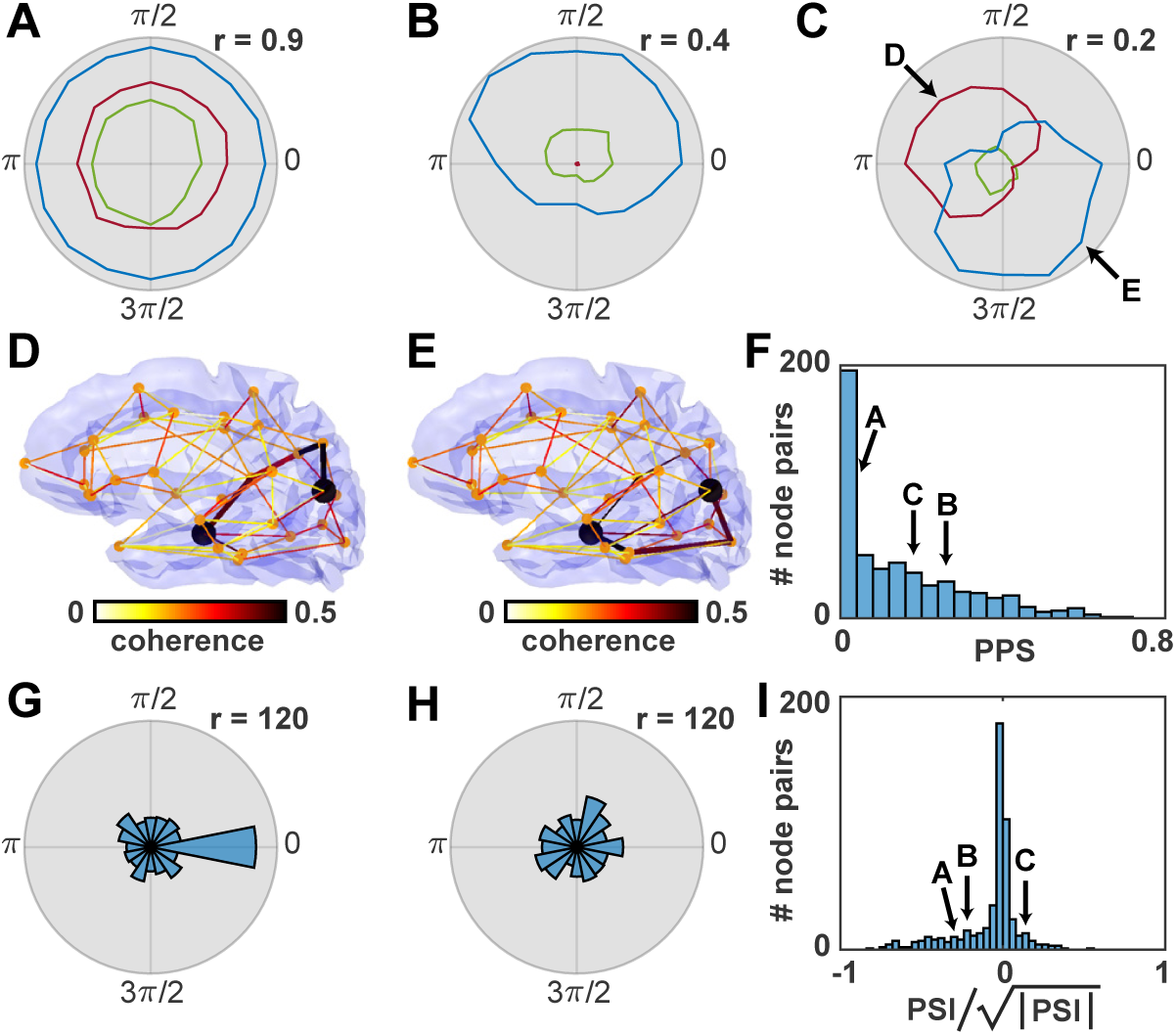
Dependence of path activation (PA) on stimulation phase offset. Panels (A-C) show the PA values (radius) for different stimulation phase offsets (angle) for example node pairs. Blue corresponds to the pathway with the strongest overall PA. Red corresponds to the second strongest pathway that has no overlapping segments with the strongest. Green curves show the strongest of the remaining pathways (with possible overlapping path segments with the former two). The two arrows in (C) indicate the phase offsets which are used in panels (D-E). (D) Connectome pathways for stimulation of the node pair shown in (C) at phase offset 1.5*π*. The two stimulated nodes are shown as two black dots. The most active pathway at this stimulation phase offset is highlighted with a stronger line width. All colors correspond to the coherence of nearest neighbours in the connection graph. (E) Similar to (D) but for stimulation phase offset 0.5*π* with a different most active pathway. (F) Histogram of pathway phase selectivity. The values of the strongest paths of examples shown in (A-C) are marked with arrows. (G) Histogram of phase differences between the stimulation phase offset where the most active pathway has the highest pathway activation and the stimulation phase offset where the coherence between the stimulated nodes was highest. (H) Similar to (G) but for the second most active pathway (excluding path overlaps with the most active pathway). (I) Histogram of normalized pathway switching index between the strongest and second strongest pathways. The values of the node pairs of examples shown in (A-C) are marked with arrows. The square-root normalization of the PSI transforms back from the space of multiplied coherence values to the original non-squared coherence space (analogous to a transformation from variance to standard-deviation).

There are several alternative ways to define the pathway activation. A similar analysis like in Fig 7 was performed with a definition of pathway activation that is weighted with the average connection strength along the path and is shown in supporting material S2 Fig. A second alternative is to calculate the minimum of all coherence values along the path instead of the product of coherence values. In this way, the communication through a pathway is only constrained by the path segment with the lowest coherence. The results using this definition are shown in supporting material S3 Fig. Both alternative definitions of the pathway activation did not change the overall results in a qualitative way, meaning that using any of these alternative definitions there are always all three types of node pairs in the network: 1) pairs with no phase selectivity in their strongest pathway (like in Fig 7A), 2) pairs with phase selectivity without switching (like in Fig 7B), 3) pairs with pathway switching (like in Fig 7C).

To further analyze how the communication via specific pathways depends on the stimulation phase offset, we define the pathway phase selectivity (PPS) of a pathway *P*_1_ similar to the PSF as

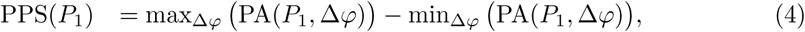

Pathways with relatively constant PA values for all stimulation phase offsets have a low PPS (example in Fig 7A), while pathways with a high variation in the PA values have a high PPS (example in Fig 7B and 7C). In general, the metric is bound to the interval between 0 and 1. The evaluation of PPS values for all node pairs results in the distribution shown in Fig 7F. The activation of pathways with a PPS close to 0 is very hard to influence with phase offsets between the nodes at their ends. However, the further the PPS grows towards 1, the more sensitive the PA is towards changes in these phase offsets. As indicated by the long tail of the PPS distribution, a significant amount of pathways in the connectome model show such phase preferences.

In the next step, we analyzed the relationship of pathway-specific phase preferences (as shown in Fig 7A-C) to the phase preferences of the stimulated nodes (as shown in Fig 5C-D). We chose the most active pathway per node pair (averaged over all stimulation phase offsets) and calculated the phase difference between the stimulation phase offset with the highest coherence between the stimulated nodes and the stimulation phase offset with the highest PA. The histogram of these phase differences is significantly different from uniform, *χ*^2^(15, *N* = 514) = 155.31, *p* < .001, and has a peak at 0 (Fig 7G). A similar analysis for the second strongest pathway (excluding all pathways with overlapping sections with the strongest path), results in a histogram that differs only slightly from a uniform distribution, *χ*^2^(15, *N* = 514) = 26.82, *p* = 0.03 (Fig 7H). Hence, we found that the pathway with the strongest PA shows a similar phase preference as the coherence between the two stimulated nodes. We take this as evidence that the interaction between the nodes through the network is strongly modulated by this pathway.

Finally, we quantified the process of switching between the strongest and second strongest pathway per node pair. To this end, we define the pathway switching index (PSI) between pathways *P*_1_ and *P*_2_ as

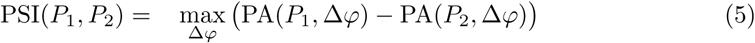

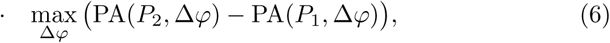

The PSI is positive if the two pathways switch their activation depending on the stimulation phase offset, meaning that at one phase offset the first path is more active and at another phase offset the second path is more active. We found that 33% (170 of 514) of node pairs have a positive PSI between their non-overlapping strongest and second strongest pathways (Fig 7I). These results suggest that in this network of 33 nodes of the human connectome many node pairs have the capacity to switch their communication between at least two different pathways with a PA characteristic similar to the example shown in Fig 7C.

## Discussion

We have carried out a computational study of cortico-cortical synchronization processes that strongly emphasizes the role of phase relationships for dynamic switches in communication pathways. In this process, we introduced a novel method to detect network interactions between pairs of cortical regions via an extrinsic stimulation scheme. Using our method, we were able to quantify the influence of different pathways on cortico-cortical coupling between all pairs of 33 brain regions. We could further identify the pathways those region pairs use to interact with each other. These pathways represent communication channels with distinct preferred interaction time lags.

One of the main findings of our work is that the pathway activation depends on the phase lag between the nodes they connect. This finding is in line with the communication-through-coherence theory that predicts neural communication to depend on oscillatory phase differences and has received support from various experimental results [20]. In a transcranial magnetic stimulation study, Elswijk and colleagues demonstrated the effect of the stimulation on primary motor cortex to depend on the oscillatory phase of the latter [47]. Similarly, Helfrich and colleagues found the performance in a visual detection task to be modulated by the phase of a 10 Hz transcranial alternating current stimulation applied to the parieto-occipital cortex [48]. Together, these experimental results support the idea of the communication-through-coherence theory that the effectiveness of neural communication between a source and a target population is modulated by the oscillatory phase of the latter. In a study using multielectrode recordings in frontal eye field and area V4 of monkeys, Gregoriou and colleagues were able to show that neural populations at both recording sites synchronized at characteristic time lags when processing a stimulus in their joint receptive field [49]. Based on the communication-through-coherence theory this may be explained by the axonal and synaptic delays of the communication pathways between these two areas, which render synchronization at this time lag most effective. The work presented in this article contributes to this notion via its systematic investigation of the dependence between pathway delays and neural synchronization. Our observation that the synchronization of two brain regions via simultaneous extrinsic stimulation has a preference for particular phase offsets between the two stimulation signals demonstrates the mechanistic role of phase lags for neural communication within a computational model. We were able to extend this finding to individual communication pathways, i.e., we showed that these pathways can express more or less strong preferences for the phase lag between the brain regions they connect. Together with the results of our toy-model simulations, which showed a direct dependence of neural synchronization on propagation delays in various connectivity motifs, this emphasizes the importance of connection delays for pathway switching. Moreover, it suggests that the effectiveness of information exchange via certain pathways depends on the phase relationship between the communicating sites.

The functional significance of these phase lag preferences is underlined by our finding that different pathways may be employed at different stimulation phase offsets between the communicating regions. Exploiting this mechanism, brain regions could dynamically switch between communication pathways by alterations in their phase lags. This flexibility in the choice of pathways could reflect a major self-organization mechanism of the brain. Specifically, the processing of transient sensory events may trigger strong phase shifts at certain sites in the brain that render certain pathways inaccessible due to their inherent delays. Our results suggest that in such a situation, the brain could employ a different set of routes to still enable communication between populations that previously relied on the now inaccessible pathway. As suggested in [12, 50], flexible switching between different processing pathways could also allow for the dynamic binding of remote neural representations into different combined representations. Both external stimuli and network intrinsic signals could act as phase-resetting mechanisms at the communicating sites and thus switch from one pathway to another within a few oscillatory cycles [51].

In our study, we considered the case of simultaneous, sinusoidal stimulation of two network nodes with a relative phase offset between the stimulation signals. In the following, we discuss how such a phase offset could arise in a biological network without external stimulation. On the one hand, a network internal stimulation could originate from a common neural driver that projects to several target sites with slightly different delays. Our results suggest that such delays could favour some communication pathways between these target sites over others. Thus, the common neural driver would not only be able to project its information to its target sites, but also choose certain pathways via which information about the input is processed. Given different neural drivers that project to the same target sites with distinct delays, different routes for integrating the information could be associated with the driving delays. On the other hand, the stimulation could also originate from an external sensory stimulus. In this scenario, different features of the stimulus are processed along different neural pathways that will converge along the sensory hierarchy. Our results suggest that the convergence of these pathways is sensitive to differences in the processing delays across pathways. Differences in processing delays could for example arise from variations in the prominence of certain stimulus features for which a certain pathway is sensitive. Depending on the prominence of the feature, the neural responses would be more or less synchronized which directly translates to differences in the processing delay further up the processing hierarchy [21]. Alternatively, the feature prominence could affect the processing delay by advancing the phase of an oscillation, by increasing the oscillation frequency, or by reducing the time to entrain other oscillators [20, 21]. This has for example been demonstrated for neural signal transmission between primary and secondary visual cortex in macaque monkeys [52]. Along these lines, Voloh and Womelsdorf provide a convincing overview with respect to different experimental scenarios in which stimulus-triggered phase resets lead to re-organizations of functional networks [53].

Different scenarios could be imagined in which input-dependent changes in the phase of a specific node affect the pattern of active output pathways of that node. One example would be the scenario described in [19, 20], where two bottom up neural drivers encoding different stimuli try to entrain a target population at different phase lags. While our results suggest that entrainment at different phase lags could affect the pathways along which the winning stimulus is processed further, this scenario of competing input stimuli is not explicitly considered in this study. Thus, future work could investigate pathway switching due to transient inputs delivered at different phases of the ongoing oscillations. To this end, a major challenge will be the required online read-out of the phase of the source node to correctly time the input. This scenario would allow to further test our proposal of dynamic switching of communication pathways in both computational models and experimental setups. A potential candidate for the latter would be brain stimulation studies, employing for example transcranial stimulation or optogenetics in combination with online recordings of neural activity [54, 55]. For a review on how such phase shifts can be induced in models of single cells and networks of interconnected cells, see [54].

We would like to propose two further future directions along which our line of research could be extended and at the same time, address two important simplifications in our model. First, we only considered discrete delays in our model, resembling average lengths of DTI-tractography based cortico-cortical tracts. However, from experimental studies it is known that long-range interactions in cortex express less sharp delay profiles [56]. Indeed, computational studies have found that the variance of delay distributions affects synchronization properties of delay-coupled systems in various ways [57, 58]. Thus, a systematic study of how stimulus phase-dependent effects on cortical routing (as reported in this study) may change with the variance of distributed delays could shed light on the validity of our results for biologically more realistic delay-coupling profiles.

Another simplification of our study concerns the Jansen-Rit model we used to represent the cortical dynamics at each of our network nodes. This model was developed to describe dynamics of average membrane potential fluctuations in three interconnected cell populations. However, the model equations have not been derived from a single cell representation of such a network [36]. As such, the model provides limited information about the underlying single cell states, such as the within-population synchronization degrees. This means that the model does not allow investigating the dependence of its synchronization with other network nodes on the degree of neural synchronization within its underlying neural populations. However, how strongly the excitability of a population varies within its oscillatory cycle clearly depends on this degree of within-population synchronization [20]. Recently developed mathematical descriptions of the macroscopic dynamics of networks of spiking neurons have overcome these problems and allow deriving the synchronization of the neural population directly from its mean-field description [59]. Unfortunately, models of cortical microcircuits have yet to be developed for these novel mean-field descriptions. With such models at hand, it would be feasible to investigate interactions between within- and between-population synchronization processes at a macroscopic scale and to investigate whether communication pathways in delay-connected networks express similar phase preference profiles as we found in this study.

Taken together, our results suggest a potential mechanism the human brain might have developed to use the physiological constraints imposed by coupling delays to its computational advantage. We have laid out several future directions of research that could help to advance and confirm these findings. From an experimental point of view, the stimulation method applied in our computational model could guide the development of brain stimulation protocols to probe dynamic switching between communication pathways in the brain. Additionally, our stimulation method and graph metrics are applicable in future theoretical studies characterizing the dynamic properties of network graphs.

## Materials and Methods

In the following, we present the detailed parameters of the neural mass model, the pre-processing of the structural connectivity, and the functional connectivity.

### Neural Mass Model

In the Jansen-Rit model, signal transmission between cell populations is realized via a convolution of average pre-synaptic firing rates with a post-synaptic response kernel. These convolution operations are mathematically described by the coupled ordinary differential equations in Fig 2 (each line describes a synaptic convolution operation) and turn pre-synaptic firing rates into post-synaptic potentials. The simple exponential form of the response kernel is in line with empirically measured post-synaptic responses [60] and provides a sufficient approximation of synaptic response time scales for our purposes. To translate post-synaptic responses back into firing rates, an instantaneous sigmoidal transform is used as shown in Fig 2 which introduces non-linearity to the model behavior.

The parameters of the neural mass model are shown in Table 2.

**Table 2.** Jansen-Rit Model Parameters.

| Param. | Value | Interpretation |
| --- | --- | --- |
| $H_e$ | 3.25 mV | avg. gain of excitatory synapses |
| $H_i$ | 22 mV | avg. gain of inhibitory synapses |
| $\tau_e$ | 10 ms | lumped time constant of excitatory synapses |
| $\tau_i$ | 20 ms | lumped time constant of inhibitory synapses |
| $c_1$ | 135 | avg. number of contacts from pyramidal cells to exc. interneurons |
| $c_2$ | $0.8 \cdot c_1$ | avg. number of contacts from exc. interneurons to pyramidal cells |
| $c_3$ | $0.25 \cdot c_1$ | avg. number of contacts from pyramidal cells to inh. interneurons |
| $c_4$ | $0.25 \cdot c_1$ | avg. number of contacts from inh. interneurons to pyramidal cells |
| $e_0$ | $2.5 \frac{\text{spikes}}{s}$ | maximum scaling of the synaptic gain |
| $r$ | $0.56 \text{ mV}^{-1}$ | steepness of the sigmoid function |
| $V_0$ | 6 mV | value with 50% of max. firing rate |
| $u_j$ | 120 - 320 $\frac{\text{spikes}}{s}$ | sub-cortical noise distributed uniformly |

They reflect the standard parametrization proposed by Jansen and Rit for modeling waxing-and-waning cortical alpha oscillations at around 10 Hz [36]. While some of those parameters (such as the connection strength ratios) were informed by experimental findings, others can be considered free parameters in bio-physiologically plausible ranges. The dynamic dependence of the Jansen-Rit model on its parameters has been investigated in various theoretical studies [36, 61–63]. For our study, it is important that each network node is kept in an oscillating dynamic regime in which the oscillation amplitude is sensitive to the network input. Since the model has several control parameters that can lead to aperiodic network behavior (see [36, 62, 63]), we decided to stick with the standard parametrization proposed by Jansen and Rit (Table 2) for each node in our network.

Thus, the only model parameter that is varied in our simulations is the average input to the model arising from the background noise, network interactions and external stimulation. As shown by Spiegler and colleagues, the Jansen-Rit model undergoes various bifurcations due to alterations of the input strength that can drive the model into bi-stable periodic or aperiodic regimes [63]. To investigate whether such undesired states could be entered under the conditions imposed by our simulations, we calculated the minimum and maximum of the average input a single node may receive in our simulations. Besides the extrinsic stimulation signal, which is centered around 0 mV, there are two rate-based inputs at each node. (1) The sub-cortical noise is modeled by a Gaussian process with a mean of 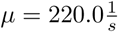. (2) The network input is ranging between 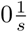 and 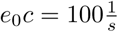. Feeding (1) and (2) into a single excitatory synapse with efficacy *H* = 3.25*mV* and *τ* = 0.01*s*, we find that it produces a synaptic input ranging between 7.5*mV* and 10*mV*. As shown by [63] the Jansen-Rit model expresses a mono-stable dynamic regime in this input range with a limit cycle being the only stable equilibrium. Variation of the input strength within this range mainly results in a modulation of the oscillation amplitude. Thus, the models considered in this study can conceptually be viewed as networks of (sparsely) coupled non-linear oscillators with discrete time delays.

If not reported otherwise, simulation results reported for those models refer to 16 minutes of simulated network behavior. We solved the model equations using the Runge-Kutta algorithm of 4th order (RK4) with an integration step-size of 1 ms. An analysis of the influence of the simulation step size on the accuracy of the simulation can be found in supporting material S1 Fig.

### Structural Connectivity and Distance Estimates

In the first step to building a bottom-up model of cortical activity, we needed to approximate the structural connections between different brain regions. As mentioned in the introduction, this can be done via DTI recordings. However, there are several technical limitations of the extent to which human SC can be approximated based on DTI, one of them being the systematic underestimation of inter-hemispheric connections [64, 65]. Thus we decided to restrict our analysis to the cerebral cortex of a single hemisphere. To this end, we used the same structural imaging data, pre-processing and probabilistic tracking pipeline as reported by Finger et al. [30], but restricted subsequent processing to the 33 regions of interest (ROIs) of the left hemisphere. This data set included diffusion- and T1-weighted images acquired from 17 healthy subjects (7 female, age mean ± s.d. = 65.6y ± 10.9y) with a 3 Tesla Siemens Skyra MRI scanner (Siemens, Erlangen, Germany) and a 32-channel head coil. The 33 ROIs were registered individually for each subject based on the ‘Desikan-Killiany’ cortical atlas available in the Freesurfer toolbox (surfer.nmr.mgh.harvard.edu) [66]. The incoming connections to each region were normalized such that they summed up to 1. Since we were only interested in synchronization along indirect pathways, we needed some connections in our model to be strictly 0. Otherwise, it would be difficult to exclude potential synchronization along very weak direct connections. Hence, we chose to set all connections below a strength of 0.1 to zero. Afterwards, we re-normalized the input to each region such that they summed up to 1. The resulting SC matrix as well as the pair-wise distances calculated from the average tract lengths are visualized in Fig 4A and 4B.

### Empirical Resting-State Functional Connectivity

Based on those SC and distance information we aimed to build a model of cortical activity able to reflect empirically observed synchronization behavior. Thus we needed empirical observations of cortical activity to evaluate our model. For this purpose, we acquired EEG data from the same 17 subjects as described above. This was done with 63 cephalic active surface electrodes arranged according to the 10/10 system (actiCAP R Brain Products GmbH, Gilching, Germany) for eight minutes of eyes-open resting-state. Again, data acquisition and pre-processing followed the same procedure as reported by Finger et al. [30]. EEG time-series from the surface electrodes were projected onto the centers of the ROIs via a linear constraint minimum variance spatial beam former [67]. The resulting source-space signals were band-pass filtered at 10 Hz and turned into analytic signals using the Hilbert transform. Subsequently, functional connectivity was evaluated as the coherence between all pairs of ROIs [68]. This resulted in the 33 x 33 functional connectivity matrix that can be observed in Fig 4C in the main paper and served as optimization target for our model.

## Supporting information

**S1 Fig.**
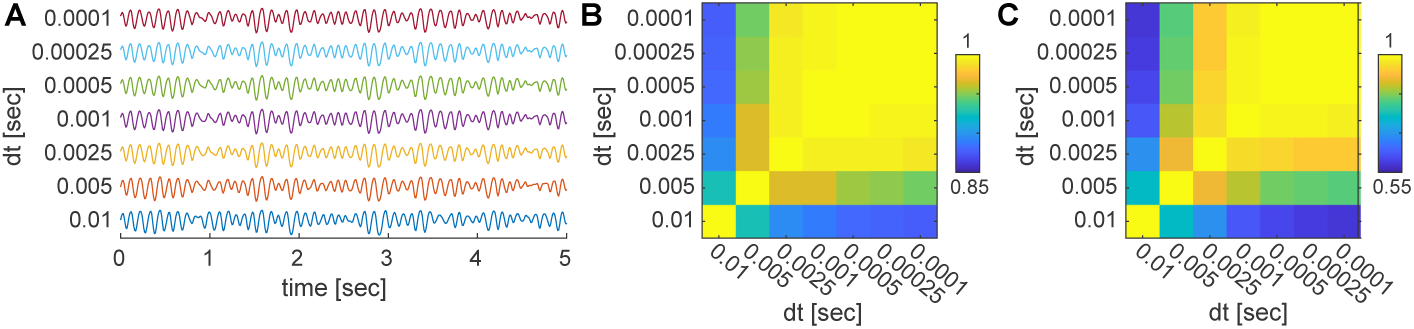
Evaluation of simulation step size. (A) comparison of the timeseries for different step sizes of the Runge-Kutta integration method. All time series are of the same node that is embedded in the connectome network of 33 nodes. (B) The correlation matrix between the timeseries using different simulation step sizes. (C) The correlation between functional connectivity matrices obtained using different simulation step sizes.

**S2 Fig.**
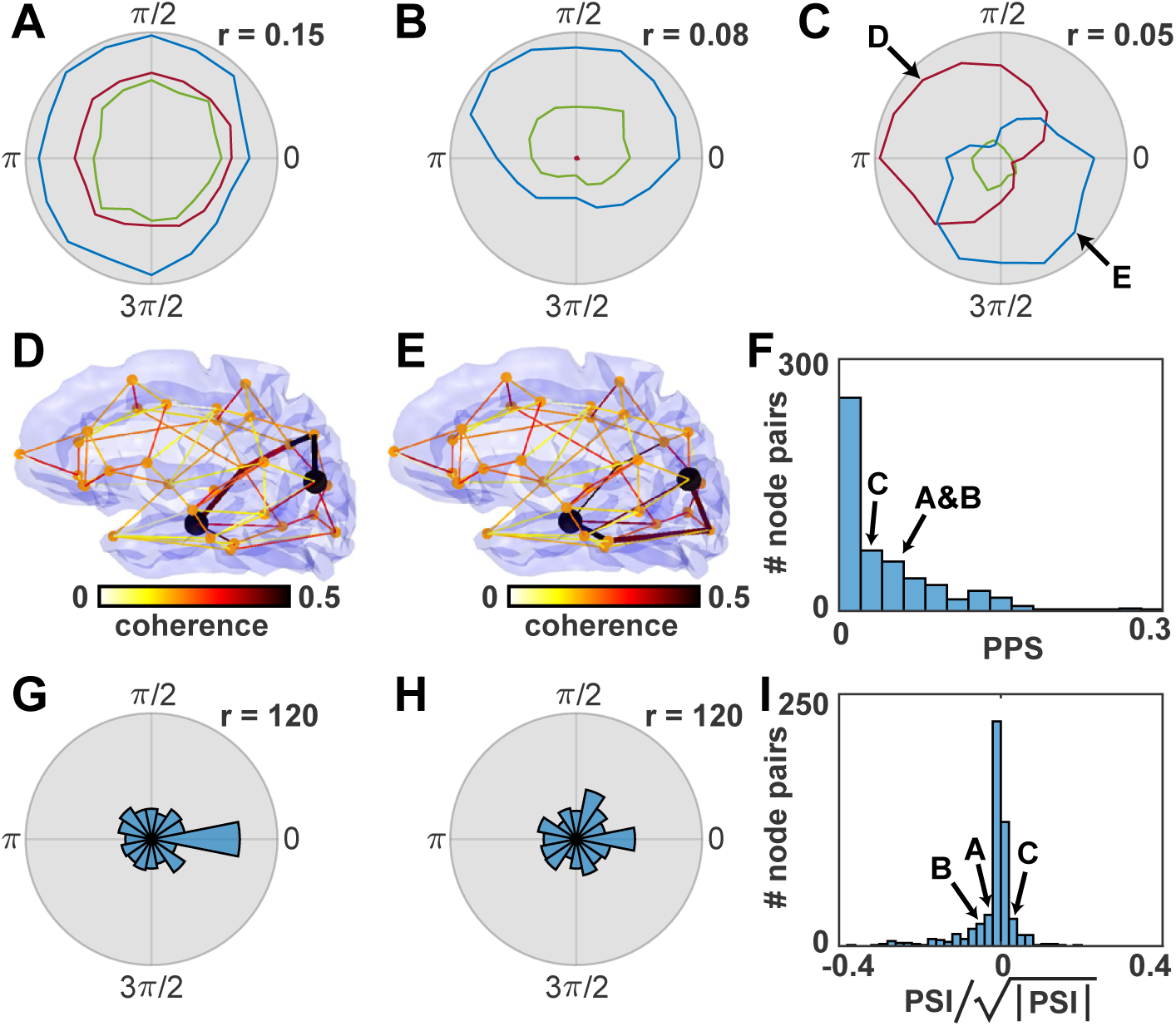
Evaluation of pathway activation with weighting of connection strengths. The panels correspond to Figure 7 in the main text. Please find the detailed panel descriptions there.

**S3 Fig.**
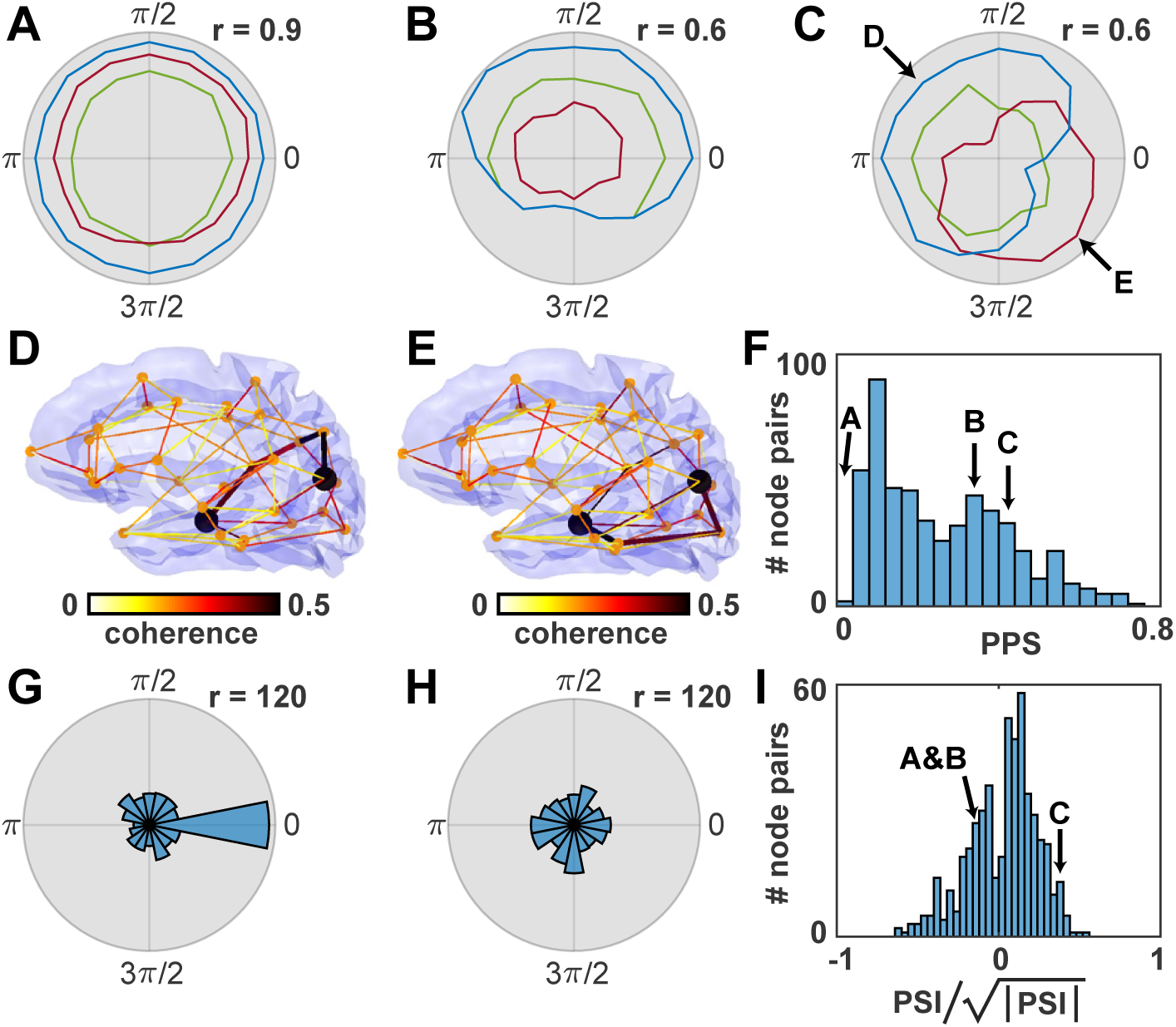
Evaluation of pathway activation using the minimum of the coherence along path segments instead of the product. The panels correspond to Figure 7 in the main text. Please find the detailed panel descriptions there.

## Notes

#### Summary of Updates

Several improvements were made based on reviewer comments.

https://github.com/hfinger/dti2eeg

https://github.com/hfinger/neuralPhasors

## References

1. Friston K, Frith C, Liddle P, Frackowiak R. Functional connectivity: the principal-component analysis of large (PET) data sets. Journal of Cerebral Blood Flow & Metabolism. 1993;13(1):5–14.

2. Biswal B, Zerrin Yetkin F, Haughton VM, Hyde JS. Functional connectivity in the motor cortex of resting human brain using echo-planar MRI. Magnetic Resonance in Medicine. 1995;34(4):537–541.

3. Damoiseaux J, Rombouts S, Barkhof F, Scheltens P, Stam C, Smith SM, et al. Consistent resting-state networks across healthy subjects. Proceedings of the National Academy of Sciences. 2006;103(37):13848–13853.

4. Van Den Heuvel MP, Pol HEH. Exploring the brain network: a review on resting-state fMRI functional connectivity. European Neuropsychopharmacology. 2010;20(8):519–534.

5. Cabral J, Kringelbach ML, Deco G. Functional connectivity dynamically evolves on multiple time-scales over a static structural connectome: Models and mechanisms. NeuroImage. 2017;.

6. Mantini D, Perrucci MG, Del Gratta C, Romani GL, Corbetta M. Electrophysiological signatures of resting state networks in the human brain. Proceedings of the National Academy of Sciences. 2007;104(32):13170–13175.

7. Brookes MJ, Hale JR, Zumer JM, Stevenson CM, Francis ST, Barnes GR, et al. Measuring functional connectivity using MEG: methodology and comparison with fcMRI. NeuroImage. 2011;56(3):1082–1104.

8. Calhoun VD, Kiehl KA, Pearlson GD. Modulation of temporally coherent brain networks estimated using ICA at rest and during cognitive tasks. Human Brain Mapping. 2008;29(7):828–838.

9. Deco G, Kringelbach ML, Jirsa VK, Ritter P. The dynamics of resting fluctuations in the brain: metastability and its dynamical cortical core. Scientific Reports. 2017;7(1):3095.

10. Greicius MD, Krasnow B, Reiss AL, Menon V. Functional connectivity in the resting brain: a network analysis of the default mode hypothesis. Proceedings of the National Academy of Sciences. 2003;100(1):253–258.

11. Fox MD, Snyder AZ, Vincent JL, Corbetta M, Van Essen DC, Raichle ME. The human brain is intrinsically organized into dynamic, anticorrelated functional networks. Proceedings of the National Academy of Sciences. 2005;102(27):9673–9678.

12. Engel AK, Gerloff C, Hilgetag CC, Nolte G. Intrinsic coupling modes: multiscale interactions in ongoing brain activity. Neuron. 2013;80(4):867–886.

13. Gross J, Schmitz F, Schnitzler I, Kessler K, Shapiro K, Hommel B, et al. Modulation of long-range neural synchrony reflects temporal limitations of visual attention in humans. Proceedings of the National Academy of Sciences. 2004;101(35):13050–13055.

14. Uddin LQ, Kelly AC, Biswal BB, Castellanos FX, Milham MP. Functional connectivity of default mode network components: correlation, anticorrelation, and causality. Human Brain Mapping. 2009;30(2):625–637.

15. Raichle ME. The brain’s default mode network. Annual Review of Neuroscience. 2015;38:433–447.

16. Klimesch W, Sauseng P, Hanslmayr S. EEG alpha oscillations: the inhibition-timing hypothesis. Brain research reviews. 2007;53(1):63–88.

17. Jensen O, Mazaheri A. Shaping functional architecture by oscillatory alpha activity: gating by inhibition. Frontiers in human neuroscience. 2010;4:186.

18. van Kerkoerle T, Self MW, Dagnino B, Gariel-Mathis MA, Poort J, van der Togt C, et al. Alpha and gamma oscillations characterize feedback and feedforward processing in monkey visual cortex. Proceedings of the National Academy of Sciences. 2014;111(40):14332. doi:10.1073/pnas.1402773111.

19. Fries P. A mechanism for cognitive dynamics: neuronal communication through neuronal coherence. Trends in Cognitive Sciences. 2005;9(10):474–480. doi:10.1016/j.tics.2005.08.011.

20. Fries P. Rhythms for Cognition: Communication through coherence. Neuron. 2015;88(1):220–235. doi:10.1016/j.neuron.2015.09.034.

21. Hahn G, Ponce-Alvarez A, Deco G, Aertsen A, Kumar A. Portraits of communication in neuronal networks. Nature Reviews Neuroscience. 2018; p. 1.

22. Engel AK, König P, Kreiter AK, Schillen TB, Singer W. Temporal coding in the visual cortex: new vistas on integration in the nervous system. Trends in Neurosciences. 1992;15(6):218–226.

23. Feng J, Jirsa VK, Ding M. Synchronization in networks with random interactions: theory and applications. Chaos: An Interdisciplinary Journal of Nonlinear Science. 2006;16(1):015109. doi:10.1063/1.2180690.

24. Deco G, Corbetta M. The dynamical balance of the brain at rest. The Neuroscientist. 2011;17(1):107–123.

25. Deco G, Jirsa VK, McIntosh AR. Emerging concepts for the dynamical organization of resting-state activity in the brain. Nature Reviews Neuroscience. 2011;12(1):43.

26. Cabral J, Hugues E, Sporns O, Deco G. Role of local network oscillations in resting-state functional connectivity. NeuroImage. 2011;57(1):130–139.

27. Rho YA, McIntosh RA, Jirsa VK. Synchrony of two brain regions predicts the blood oxygen level dependent activity of a third. Brain Connectivity. 2011;1(1):73–80. doi:10.1089/brain.2011.0009.

28. Siegel M, Donner TH, Engel AK. Spectral fingerprints of large-scale neuronal interactions. Nature Reviews Neuroscience. 2012;13(2):121.

29. Moon JY, Lee U, Blain-Moraes S, Mashour GA. General relationship of global topology, local dynamics, and directionality in large-scale brain networks. PLOS Computational Biology. 2015;11(4):e1004225. doi:10.1371/journal.pcbi.1004225.

30. Finger H, Bönstrup M, Cheng B, Messé A, Hilgetag C, Thomalla G, et al. Modeling of large-scale functional brain networks based on structural connectivity from DTI: comparison with EEG derived phase coupling networks and evaluation of alternative methods along the modeling path. PLoS Computational Biology. 2016;12(8):e1005025.

31. Petkoski S, Spiegler A, Proix T, Aram P, Temprado JJ, Jirsa VK. Heterogeneity of time delays determines synchronization of coupled oscillators. Physical Review E. 2016;94(1):012209. doi:10.1103/PhysRevE.94.012209.

32. Petkoski S, Palva JM, Jirsa VK. Phase-lags in large scale brain synchronization: methodological considerations and in-silico analysis. PLOS Computational Biology. 2018;14(7):e1006160. doi:10.1371/journal.pcbi.1006160.

33. Roxin A, Brunel N, Hansel D. Role of delays in shaping spatiotemsporal dynamics of neuronal activity in large networks. Physical Review Letters. 2005;94(23):238103. doi:10.1103/PhysRevLett.94.238103.

34. Roxin A, Montbrió E. How effective delays shape oscillatory dynamics in neuronal networks. Physica D: Nonlinear Phenomena. 2011;240(3):323–345. doi:10.1016/j.physd.2010.09.009.

35. Ghosh A, Rho Y, McIntosh AR, Kötter R, Jirsa VK. Noise during rest enables the exploration of the brain’s dynamic repertoire. PLOS Computational Biology. 2008;4(10):e1000196. doi:10.1371/journal.pcbi.1000196.

36. Jansen BH, Rit VG. Electroencephalogram and visual evoked potential generation in a mathematical model of coupled cortical columns. Biological Cybernetics. 1995;73(4):357–366.

37. Felleman DJ, Van Essen DC. Distributed Hierarchical Processing in the Primate Cerebral Cortex. Cerebral Cortex. 1991;1(1):1–47. doi:10.1093/cercor/1.1.1-a.

38. Sotero RC, Trujillo-Barreto NJ, Iturria-Medina Y, Carbonell F, Jimenez JC. Realistically coupled neural mass models can generate EEG rhythms. Neural Computation. 2007;19(2):478–512. doi:10.1162/neco.2007.19.2.478.

39. Du X, Jansen BH. A neural network model of normal and abnormal auditory information processing. Neural Networks. 2011;24(6):568–574. doi:10.1016/j.neunet.2011.03.002.

40. Jedynak M, Pons AJ, Garcia-Ojalvo J. Cross-frequency transfer in a stochastically driven mesoscopic neuronal model. Frontiers in Computational Neuroscience. 2015;9. doi:10.3389/fncom.2015.00014.

41. Kunze T, Hunold A, Haueisen J, Jirsa V, Spiegler A. Transcranial direct current stimulation changes resting state functional connectivity: A large-scale brain network modeling study. Transcranial electric stimulation (tES) and Neuroimaging. 2016;140(Supplement C):174–187. doi:10.1016/j.neuroimage.2016.02.015.

42. Ahmadizadeh S, Karoly PJ, Nešić D, Grayden DB, Cook MJ, Soudry D, et al. Bifurcation analysis of two coupled Jansen-Rit neural mass models. PLOS ONE. 2018;13(3):e0192842. doi:10.1371/journal.pone.0192842.

43. Vicente R, Gollo LL, Mirasso CR, Fischer I, Pipa G. Dynamical relaying can yield zero time lag neuronal synchrony despite long conduction delays. Proceedings of the National Academy of Sciences. 2008;105(44):17157–17162. doi:10.1073/pnas.0809353105.

44. König P, Schillen TB. Stimulus-dependent assembly formation of oscillatory responses: I. Synchronization. Neural Computation. 1991;3(2):155–166.

45. Messé A, Rudrauf D, Giron A, Marrelec G. Predicting functional connectivity from structural connectivity via computational models using MRI: an extensive comparison study. NeuroImage. 2015;111:65–75.

46. Boyland PL. Bifurcations of circle maps: Arnol’d tongues, bistability and rotation intervals. Communications in Mathematical Physics. 1986;106(3):353–381.

47. Elswijk Gv, Maij F, Schoffelen JM, Overeem S, Stegeman DF, Fries P. Corticospinal beta-band synchronization entails rhythmic gain modulation. Journal of Neuroscience. 2010;30(12):4481–4488. doi:10.1523/JNEUROSCI.2794-09.2010.

48. Helfrich RF, Schneider TR, Rach S, Trautmann-Lengsfeld SA, Engel AK, Herrmann CS. Entrainment of brain oscillations by transcranial alternating current stimulation. Current Biology. 2014;24(3):333–339.

49. Gregoriou GG, Gotts SJ, Zhou H, Desimone R. High-frequency, long-range coupling between prefrontal and visual cortex during attention. Science. 2009;324(5931):1207–1210. doi:10.1126/science.1171402.

50. Engel AK, Singer W. Temporal binding and the neural correlates of sensory awareness. Trends in Cognitive Sciences. 2001;5(1):16–25. doi:10.1016/S1364-6613(00)01568-0.

51. Canavier CC. Phase-resetting as a tool of information transmission. Current opinion in neurobiology. 2015;31:206–213. doi:10.1016/j.conb.2014.12.003.

52. Zandvakili A, Kohn A. Coordinated Neuronal Activity Enhances Corticocortical Communication. Neuron. 2015;87(4):827–839. doi:10.1016/j.neuron.2015.07.026.

53. Voloh B, Womelsdorf T. A Role of Phase-Resetting in Coordinating Large Scale Neural Networks During Attention and Goal-Directed Behavior. Frontiers in Systems Neuroscience. 2016;10. doi:10.3389/fnsys.2016.00018.

54. Tiesinga PHE, Sejnowski TJ. Mechanisms for phase shifting in cortical networks and their role in communication through coherence. Frontiers in Human Neuroscience. 2010;4. doi:10.3389/fnhum.2010.00196.

55. Yazdan-Shahmorad A, Silversmith DB, Kharazia V, Sabes PN. Targeted cortical reorganization using optogenetics in non-human primates. eLife. 2018;7:e31034.

56. Nunez PL, Cutillo BA. Neocortical dynamics and human EEG rhythms. Oxford University Press, USA; 1995.

57. Eckhorn R, Gail AM, Bruns A, Gabriel A, Al-Shaikhli B, Saam M. Different types of signal coupling in the visual cortex related to neural mechanisms of associative Processing and perception. IEEE Transactions on Neural Networks. 2004;15(5):1039–1052. doi:10.1109/TNN.2004.833130.

58. Atay F, Hutt A. Neural fields with distributed transmission speeds and long-range feedback delays. SIAM Journal on Applied Dynamical Systems. 2006;5(4):670–698. doi:10.1137/050629367.

59. Montbrió E, Pazó D, Roxin A. Macroscopic description for networks of spiking neurons. Physical Review X. 2015;5(2):021028. doi:10.1103/PhysRevX.5.021028.

60. Gerstner W, Kistler WM, Naud R, Paninski L. Neuronal dynamics: From single neurons to networks and models of cognition. Cambridge University Press; 2014.

61. David O, Friston KJ. A neural mass model for MEG/EEG: coupling and neuronal dynamics. NeuroImage. 2003;20(3):1743–1755.

62. Grimbert F, Faugeras O. Bifurcation analysis of Jansen’s neural mass model. Neural Computation. 2006;18(12):3052–3068. doi:10.1162/neco.2006.18.12.3052.

63. Spiegler A, Kiebel SJ, Atay FM, Knösche TR. Bifurcation analysis of neural mass models: impact of extrinsic inputs and dendritic time constants. Computational Models of the Brain. 2010;52(3):1041–1058. doi:10.1016/j.neuroimage.2009.12.081.

64. Li L, Rilling JK, Preuss TM, Glasser MF, Damen FW, Hu X. Quantitative assessment of a framework for creating anatomical brain networks via global tractography. NeuroImage. 2012;61(4):1017–1030.

65. Wedeen VJ, Wang R, Schmahmann JD, Benner T, Tseng WYI, Dai G, et al. Diffusion spectrum magnetic resonance imaging (DSI) tractography of crossing fibers. NeuroImage. 2008;41(4):1267–1277.

66. Desikan RS, Ségonne F, Fischl B, Quinn BT, Dickerson BC, Blacker D, et al. An automated labeling system for subdividing the human cerebral cortex on MRI scans into gyral based regions of interest. NeuroImage. 2006;31(3):968–980.

67. Van Veen BD, Van Drongelen W, Yuchtman M, Suzuki A. Localization of brain electrical activity via linearly constrained minimum variance spatial filtering. IEEE Transactions on Biomedical Engineering. 1997;44(9):867–880.

68. Andrew C, Pfurtscheller G. Event-related coherence as a tool for studying dynamic interaction of brain regions. Electroencephalography and Clinical Neurophysiology. 1996;98(2):144–148.

